# Self-organization of collective escape in pigeon flocks

**DOI:** 10.1101/2021.07.04.450902

**Authors:** Marina Papadopoulou, Hanno Hildenbrandt, Daniel W.E. Sankey, Steven J. Portugal, Charlotte K. Hemelrijk

## Abstract

Bird flocks under predation demonstrate complex patterns of collective escape. These patterns may emerge by self-organization from simple interactions among group-members. Computational models have been shown to be valuable for identifying the behavioral rules that may govern these interactions among individuals during collective motion. However, our knowledge of such rules for collective escape is limited by the lack of quantitative data on bird flocks under predation in the field. In the present study, we analyze the first dataset of GPS trajectories of pigeons in airborne flocks attacked by a robotic falcon in order to build a species-specific model of collective escape. We use our model to examine a recently identified distance-dependent pattern of collective behavior that shows an increase in the escape frequency of pigeons when the predator is closer. We first extract from the empirical data the characteristics of pigeon flocks regarding their shape and internal structure (bearing angle and distance to nearest neighbours). Combining these with information on their coordination from the literature, we build an agent-based model tuned to pigeons’ collective escape. We show that the pattern of increased escape frequency closer to the predator arises without flock-members prioritizing escape when the predator is near. Instead, it emerges through self-organization from an individual rule of predator-avoidance that is independent of predator-prey distance. During this self-organization process, we uncover a role of hysteresis and show that flock members increase their consensus over the escape direction and turn collectively as the predator gets closer. Our results suggest that coordination among flock-members, combined with simple escape rules, reduces the cognitive costs of tracking the predator. Such rules that are independent of predator-prey distance can now be examined in other species. Finally, we emphasize on the important role of computational models in the interpretation of empirical findings of collective behavior.

**Author summary:** Bird flocks show fascinating patterns of collective motion, particularly when escaping a predator. Little is however known about their underlying mechanisms. We fill this gap by firstly analyzing GPS data of pigeon flocks under attack by a robotic-predator and secondly, studying their collective escape in a computer simulation. Previous research on pigeons has revealed that flock-members turn away from the predator more the closer the predator gets. Using computer simulations that are based on pigeon-specific characteristics of motion and coordination among individuals, we study what escape rules at the individual level may underlie this distance-dependent pattern. We show that even if individuals do not intend to escape more when the predator is closer, their escape frequency still increases the closer they get to the predator. This happens by self-organization from the coordination among individuals and despite their tendency to turn away from the predator being constant. A key aspect of this process is the increasing consensus among flock members over the escape direction when the predator gets closer.

## Introduction

Computational models based on self-organization are a valuable tool to disentangle the processes underlying patterns of complex biological systems. Such models show that many collective patterns that are seen in nature are emergent, i.e. not represented in the behavioral rules of the group members [1]. Several examples of such emergent phenomena have been studied in models of collective motion. In simulated fish schools, the oblong group shape (a characteristic of real schools) emerges from coordinating individuals avoiding their nearest neighbor by slowing down or turning away [2, 3]. Milling (collective circular motion) emerges in a minimal model of collective motion when individuals are limited in their field of view and angular velocity [4]. In more complex models, changes between milling and schooling behavior at the group level (phase transitions) spontaneously arise from a single set of behavioral rules at the individual level [5, 6]. Realistic group shapes emerge from the specifics of individuals’ locomotion, i.e. flying versus swimming motion [3]. Swarm shapes may also depend on the preceding shape of the group, a phenomenon called hysteresis [5]. In total, these models help us to determine what behavioral rules at the individual level are essential for a collective pattern to arise.

Models of collective motion have also been used to study collective patterns of escape from a predator. Inada and Kawachi (2002) developed a two-dimensional model of fish schools under attack. Despite modeling a single rule of escape at the individual level, several collective escape patterns with high variability of group shape emerged [7]. In airborne flocks of birds, the only pattern of collective escape that has been studied in detail is the agitation wave: a dark band that moves from one side of the group to the other [8–10]. A wave was initially assumed to relate to increased density among individuals when they flee away from the predator and thus come closer to each other [11]. The individual-level rules of this process were studied in a three-dimensional model of European starlings (*Sturnus vulgaris*) [12] where a few initiators were performing an escape motion that was subsequently copied by their neighbors [13]. In the simulated flocks, a dark band was visible only during a turning escape maneuver (instead of a forward acceleration). This was because, while banking, the larger surface of the wings becomes visible to an observer. This lead to the conclusion that this band can reflect an orientation, instead of a density, wave [13]. This study highlighted the importance of such theoretical experiments in the understanding of which behavior underlie complex collective patterns.

Given across species differences in collective behavior [14–16], a computational model should be adjusted to empirical data in order to study in detail specific patterns seen in nature [12, 17–19]. Collecting these quantitative data can be challenging. For bird flocks, results of field experiments using stereo photography have been extremely valuable for validating model conclusions and formulating hypotheses [12, 14, 16, 20]. However, due to the specifics of this technique, the positions of flock members can be reconstructed only for a few seconds of flight while the flock is passing through a stationary set of cameras. This poses a limitation for studying their collective escape: capturing a full escape sequence within the narrow frame of the cameras is unlikely. Furthermore, the challenge of controlling an avian predator in the field and tracking its motion during a pursuit limits techniques that allow the collection of full trajectories of flock members, such as GPS devices [15, 21]. Due to these constrains, until recently the collective escape of airborne flocks has been empirically studied only through video footage, focusing on qualitative descriptions of the observed collective patterns [8, 10, 22]. This lack of quantitative data has limited the development of computational models of bird flocks as well as our knowledge on the underlying processes of their collective escape [13, 23].

GPS data of a flock under predation have recently been collected using a robotic falcon to attack flocks of homing pigeons (*Columba livia*) [24]. Based on the tracks of escaping individuals during a pursuit, Sankey *et al*. (2021) identified a new distance-dependent pattern of collective escape: when the predator is close, pigeons are turning away from its heading more often than they align with their flock mates. It is however not clear whether this reflects a distance-dependent behavioral rule of individual pigeons or an emergent property. The trajectories of flock members [24], along with previous findings on local interactions in pigeon flocks [15, 21], provide the necessary information to study this phenomenon in a computational model.

The aim of the present paper is to study the empirical finding that pigeons in a flock turn away from the predator more frequently when the predator is closer [24]. We first analyze the GPS trajectories of individual pigeons to define their flock shape and internal structure (bearing angle and distance to nearest neighbor) [24]. We combine these data with information on the specifics of flocking in homing pigeons [15, 21, 24] and build a realistic computational model of pigeons’ collective escape [17]. We use our model to investigate whether flock members need an individual rule to escape more at closer distances to the predator for their escape frequency to scale with predator-prey distance. We model a predator-avoidance mechanism that reflects our null hypothesis: pigeons turn to escape without taking into account their distance to the predator. We analyze the effect of predator-avoidance and coordination among flock members during an attack. We confirm our hypothesis and conclude that the frequency of escape turns increases with decreasing distance to the predator as an emergent property during collective escape.

## Materials and methods

### Flocks of pigeons under attack

#### Empirical data

We used pre-proccessed trajectories of flocks of homing pigeons (*Columba livia*) collected by Sankey *et al*. [24]. All flock members were trained to fly back to their home after being released at a site approximately 5 km away. A robotic falcon [24, 25], similar to a peregrine falcon (*Falco peregrinus*) in appearance and locomotion, was remotely controlled to attack the flocks at their release and to chase them until they leave the site. Both prey and predator were mounted with GPS devices sampling with a frequency of 0.2 seconds (see [24] for full details).

### A data-inspired computational model

We developed an agent-based model, named *HoPE* (Homing Pigeons Escape), that simulates airborne flocks of pigeons under attack by a predator. We present our models based on elements of the ODD (Overview, Design concepts, Detail) protocol [26].

#### Principles, entities, and process overview

Our model is based on self-organization [1]. It consists of pigeon- and predator-like agents. Each simulation includes several pigeon-agents (also referred to as ‘prey’) that form a flock, and one predator-agent that pursues and attacks the flock. We adjusted the coordination rules of alignment, avoidance and attraction among nearby pigeons-agents [5, 27, 28] to known behavior of pigeons [15, 21, 24]. Parameters not found in the literature were determined through calibration with empirical measurements of several flock’s characteristics, namely the distributions of individuals’ speed, nearest-neighbor distance, and relative position of nearest neighbor (bearing angle and distance). All parameters in our model are presented in Table 1.

**Table 1.** The parameters of the HoPE model. The majority of parameter values are taken from previous empirical work on pigeons flocks [15, 21, 24]. We decide the value of parameters that could not be inferred from the literature by calibration [17] through comparisons with the empirical data of Sankey et al. [24] and visual testing to ensure realistic flock formations.

| Parameter | Name - Description | Value | Unit | Reference |
| --- | --- | --- | --- | --- |
| **Collective motion** |  |  |  |  |
| $u_c$ | Mean cruise speed | 16 | $m/s$ | [21] |
| $u_c^{sd}$ | Standard deviation of cruise speed | 2 | $m/s$ | [21] |
| $m$ | Pigeons’ body mass | 0.45 | kg | [21] |
| $\theta_{FoV}$ | Angle of field of view | 215 | degrees | [15] |
| $\theta_{FoV}^f$ | Angle of frontal part of field of view | 180 | degrees | [15] |
| $n_{topo}$ | Number of interacting neighbors for coordination | 7 | individuals | [24] |
| $d_{min}^s$ | Minimum distance for separation force | 1 | m | [15] |
| $d_{min}^\eta$ | Minimum mean distance to neighbors for acceleration | 2 | m | [15] |
| $d_{max}^\eta$ | Maximum mean distance to neighbors for acceleration | 10 | m | [15] |
| $\kappa$ | Strength of deceleration | 0.5 | - | [15] |
| $w_a$ | Weighting factor of alignment force | 10 | - | - |
| $w_s$ | Weighting factor of separation force | 5 | - | - |
| $w_{ct}$ | Weighting factor of turning cohesion force | 2.5 | - | - |
| $w_{cs}$ | Weighting factor of speed-based cohesion force | 5 | - | - |
| $w_f$ | Weighting factor of flight-control force | 0.2 | - | - |
| $\beta_w$ | Range of uniform distribution for the weighting factor of the random-error force | 2 | - | - |
| **Collective escape** |  |  |  |  |
| $w_p$ | Weighting factor of pigeons’ predator avoidance | 2 | - | - |
| $d_{max}^e$ | Minimum distance for pigeons’ predator avoidance | 50 | m | [24] |
| $\zeta_p$ | Predator’s pursuit speed relative to the flock’s | 1.0 | - | - |
| $\xi_p$ | Predator’s attack speed relative to the flock’s | 1.5 | - | - |
| $d_h$ | Distance of predator’s pursuit | 30 - 40 | m | [24] |
| $\theta_p$ | Bearing angle of predator’s pursuit | 130 - 225 | degrees | - |
| $T_h$ | Predator’s pursuit duration | 30 | s | - |
| $T_a$ | Predator’s attack duration | 20 | s | - |

Table 1. Continued
| Parameter | Name - Description | Value | Unit | Reference |
| --- | --- | --- | --- | --- |
| **Simulation** |  |  |  |  |
| $dt$ | Integration time | 0.005 | s | - |
| $\Delta t$ | Update frequency | 0.02 | s | - |
|  | Sampling frequency | 0.2 | s | [24] |

#### Motion and sensing

Our agents move in a large two-dimensional open space. They are defined by a position vector 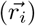 in the global space, and a velocity (*v_i_*) and a heading vector (*ĥ_i_*) in their own coordinate system. We assume that agents always have their heading in the direction of their velocity (non-slip assumption). Each agent senses the position and heading of other agents in its field of view (controlled by the angle *θ_FoV_*). The field of view of pigeon-agents is set to 215°, and is divided into a front (±90° around their heading, 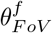) and a side area [15].

From all sensed individuals, each agent only interacts with a fixed number of closest neighbors (referred to as ‘topological neighbors’, *n_topo_*) [12, 29]. Sankey *et al*. 2021 [24] estimated that the topological range of interaction may differ between small and large flocks, and between alignment and centroid-attraction. Their method is, however, not well established and was unable to reveal the (true) topological range used in our model (S4 Fig). Because of this, and aiming to reduce the complexity of our model [17], we chose a constant number of 7 topological neighbors for both alignment and attraction across different flock sizes [16, 29].

The motion of pigeon-agents in our model is dictated by an external pseudo-force 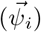, which is the sum of a coordination force, an escape force, and an internal flight control system (Fig 1). Since birds cannot perfectly collect and respond to information about their surroundings (e.g. average position and direction of neighbors), we included a random error in their motion. Specifically, a noise scalar (*ϵ_i_*) is randomly sampled by a uniform distribution (with a range *β_w_*) and multiplied with a unit vector perpendicular to the agent’s headings, forming a pseudo-force that affects the agents’ turning motion 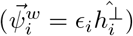.

**Fig 1.**
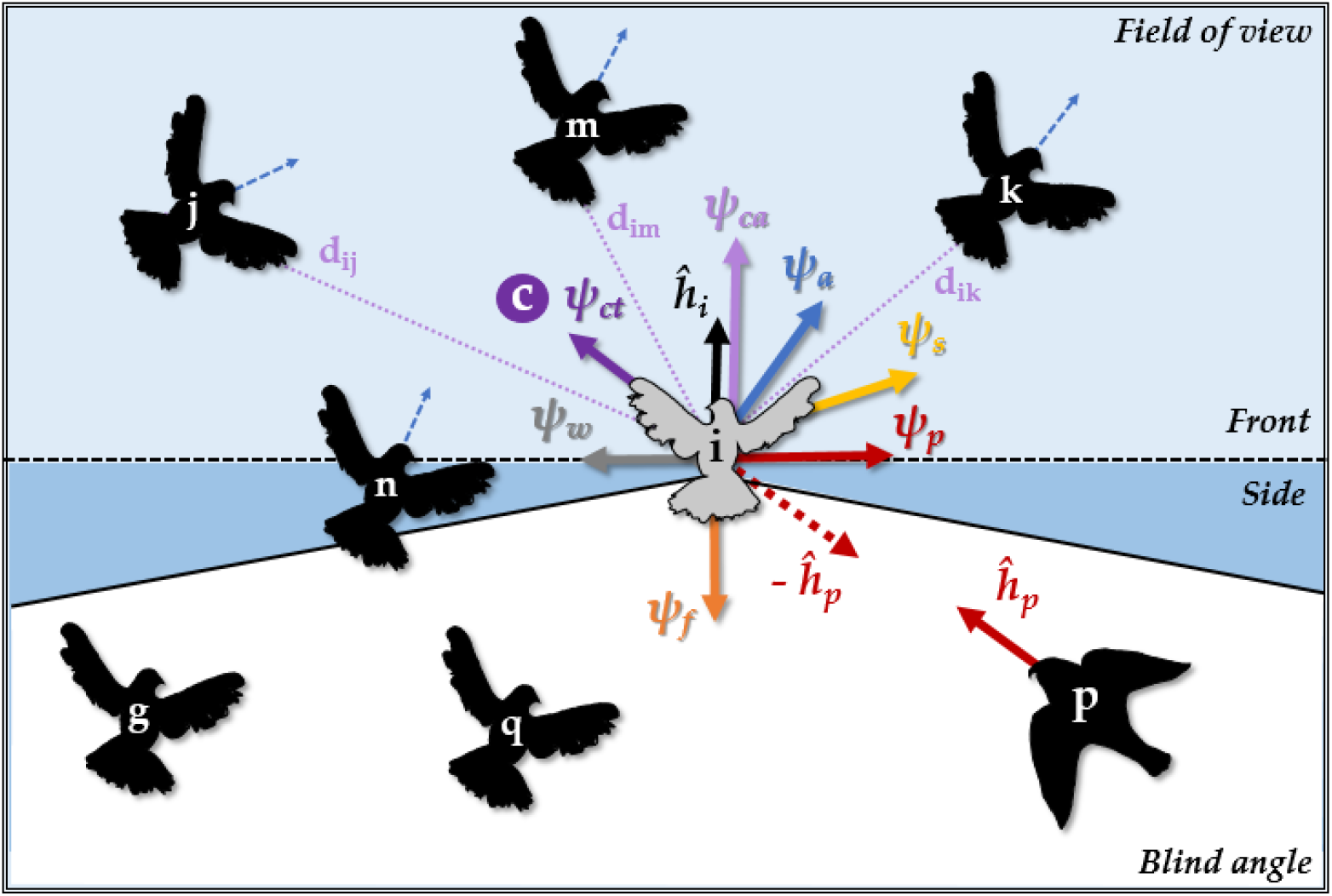
Collective motion schematic in the computational model *HoPE*. The colored areas represent the field of view of a focal individual *i* with heading *ĥ_i_*, split into a ‘front’ (light blue, 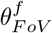) and ‘side’ area (blue). Based on its total field of view (*θ_FoV_*), *i* interacts with its topological neighbors *n*,*j*, *m*, and *k*. Agents *g* and *q* are in the blind angle of *i* and are thus ignored. All pseudo-forces (*ψ*) acting on *i* are represented by colored arrows. Alignment (*ψ_a_*) is the average vector of topological neighbors’ headings (indicated by blue doted arrows). Centroid attraction (*ψ_ct_*) is the vector from i’s position to the center of the topological neighborhood (*c*). Accelerating attraction (*ψ_ca_*) is a vector in the direction of motion (aligned with heading *ĥ_i_*), depending on the distances of *i* to the neighbors in its front field of view, namely *j*, *m* and *k* (with distances *d_ij_*, *d_im_* and *d_ik_* respectively). If there are no agents in the front field of view, then *ψ_ca_* is negative. Separation (*ψ_s_*) is the vector away from the position of the nearest neighbor n. Avoidance of the predator (*ψ_p_*) is a perpendicular vector in the direction away from the predator’s heading (*ĥ_p_*). Vector *ψ_f_* represents the flight-control force that drags i towards its preferred speed. *ψ_w_* is the force to create random-error in the orientation of the agents.

#### Individual variation and speed control

Several models of collective behavior assume constant and often identical speed of all group members [4–6, 19]. Homing pigeons, however, have a preferred ‘solo’ speed of flight, from which they deviate by up to 2 *m*/*s* when flying in a flock [21]. When flying in pairs, they accelerate to catch up with their frontal neighbor (the further away the neighbor the higher the acceleration) and slow down if they are in front [15]. According to these findings, agents in our model have different preferred speeds from which they can deviate to stay with their flock mates.

At initialization, the model randomly samples a preferred speed for each agent from a uniform distribution with interval length of 4 *m*/*s* (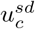 around the mean *u_c_*, Table 1). To reflect the inability of individuals to deviate from their preferred speed for a prolonged period [21], we modeled a drag force 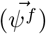 that pulls agents back to it (Eq 1). This force increases with increasing deviation from the preferred flight speed, according to:

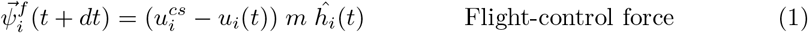

where 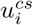 is the personalised speed of agent *i* with mass *m*, and *u_i_*(*t*) is its speed on timestep *t*.

#### Coordination

Flock formation emerges from simple rules of among-individuals coordination: attraction, avoidance and alignment [5, 12, 27, 28]. In our model, we parameterized alignment to be the strongest among our steering forces, as found in homing pigeons [24]. The alignment force 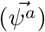 has the direction of the average heading of all topological neighbors. Centroid-attraction in our model is relatively weak, given that coherence in flocks of pigeons is mediated mostly by speed adjustment [15, 24]. We thus introduced a ‘speed-attraction’ force 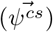: individuals accelerate if they have neighbors within their front field of view and decelerate if they do not sense any individuals nearby (their field of view is empty). The strength of this acceleration force increases with increasing average distance to all frontal neighbors, according to a smootherstep function [30] from 2 to 10 m (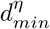 and 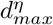) based on [15]. Deceleration is constant to the half of acceleration’s maximum (based on the scaling factor *κ*) [15]. The centroid-attraction force 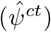 is the unit vector with direction from the position of the focal individual to the average position of its topological neighbors.

Lastly, separation among pigeons is mediated by turning when neighbors are within a 1-meter distance from each other [15]. Similarly in our model, an avoidance force 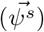 pushes agents to turn away from the position of their closest neighbor (instead of all topological neighbors) if they are too close (according to the minimum-separation distance, 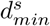). We parameterise this switch from 7 to 1 topological range for separation to increase the resemblance of our model to empirical data (following previous theoretical work of [20]), and since there is no, to our knowledge, previous research on the real topological range for separation in pigeon flocks.

To balance these forces according to real pigeons [15, 24], we implemented a weighted sum (Eq 2) with factors: 1 for the centroid-attraction weight (*w_ct_*), −1 to 2 for the speed-attraction weight (*w_cs_*), 2 for the avoidance weight (*w_s_*), and 4 for the alignment weight (*w_a_*). Their exact values are established during calibration (Table 1). The total coordination force is calculated by:

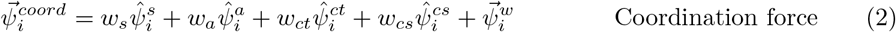

#### Escape motion

According to the empirical findings, the heading of the robotic falcon had a larger effect on the turn-away motion of pigeons than its position [24]. Hence, our pigeon-agents avoid the predator based on an escape force 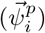 that points away from the predator’s heading. The magnitude of this force (*w_p_*) is constant and independent from the predator-prey distance:

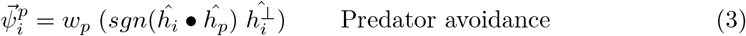

where *ĥ_p_* is the heading of the predator-agent, and *ĥ_i_* • *ĥ_p_* its dot product with the pigeon-agent’s heading. Pigeon-agent do not sense the predator’s position and thus have no information about their distance to it.

#### The predator

We model our predator-agents to resemble the motion of the robotic falcon [24] in order for our results to be comparable with the empirical data. Each simulation includes only one predator. A few seconds after the flock is formed, the predator is positioned at a given distance behind it (*d_h_*). The predator-agent follows the position of the pigeon-agent closest to it, with the speed as the prey (based on the scaling factor *ζ_p_*) from an bearing angle *θ_p_* relative to the flock’s heading. After some time (*T_h_*), the predator-agent will speed up attacking its target (with a speed scaling from the target’s speed, 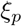) with a random error added to its motion. The predator’s target is selected based on two alternative strategies: the target is the pigeon-agent that is closer to the predator at every time step during the attack (‘chase’ strategy) or the closest pigeon-agent at the beginning of the attack (‘lock-on’ strategy). After an attack (*T_α_*), the predator is automatically re-positioned far away from the flock.

#### Update and integration

We use two different timescales for update and integration in our model, following previous biologically-relevant computational models of collective motion [12, 31]. During update steps (*t* + Δ*t*), agents collect information from their environment and the pseudo-forces acting on them are recalculated. All pigeon-agents update their information asynchronously, but with the same frequency (Δ*t*). At each integration step (*dt*), the total force for each agent is composed from:

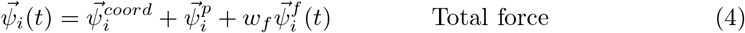

where *w_f_* is the calibrated weight for cruise speed control, and 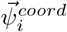 and 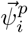 are the pseudo-forces for coordination (Eq 2) and predator-avoidance (Eq 3) calculated at the last update step. Based on this force, the acceleration, velocity, and position of all agents is updated at each integration step according to Newton’s laws of motion and using Euler’s integration (specifically the midpoint method [32]). The new heading of each agent is then the normalised form of its new velocity:

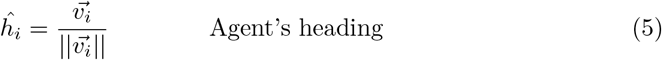

### Experiments

For our main analysis, to test the effect of predator-prey distance on the pigeons’ escape frequency, we performed two types of experiments. In both experiments, the predator is using the ‘chase’ strategy. First, we run simulations in which prey is not reacting to the predator. These simulated data were used as control. In the second experiment, pigeon-agents avoid the predator-agent without accounting for its distance to them (as described in Eq 3). We ran 1000 simulations (hunting cycles) for each experiment, while varying the direction of the predator’s attack (from the left, the middle, and the right side behind the flock relative to the flight direction, *θ_p_*) and the flock size (8, 10, 27, and 34 individuals, as in the field experiment of homing pigeons [24]). For our analysis, we combined the results of all simulations into one dataset per experiment.

To test the effect of the predator’s strategy, we repeated our main experiments using (1) the ‘lock-on’ strategy and (2) reverted versions of the two strategies. Specifically, the predator was placed automatically in close proximity to the flock (5 m) and given a speed lower than its targets’ speed; this resulted in the predator performing approximately the opposite motion of a real attack. We performed this ‘reverted’ experiments to test effects of hysteresis [5] during a collective escape.

### Analysis

The first seconds of each simulated flight are discarded to avoid effects of initial conditions. Similarly, the taking-off part of the real trajectories was excluded from our analysis. For the analysis on the predator-prey distance, data of individuals that are more than 60 meters [24] away from the predator are also discarded.

#### Defining a flock of pigeons

For both empirical and simulated data, we extracted the time-series of individual speed and nearest neighbor distance. We further estimated the bearing angle between every focal individual and each one of its neighbors based on its heading and the line connecting their positions. The shape of each flock was calculated based on the minimum volume bounding-box method [12, 33]. Specifically, we drew a bounding box of minimum area that included all flock-members’ positions, with axes aligned to the output of a principal component analysis of their coordinates (method used in [12]). To use as a proxy of flock shape, we estimated the angle between the shortest side (dimension) of this box and the heading vector of the flock (average of all flock-members) [12]. The closer this angle is to 0 degrees, the more perfectly ‘wide’ the flock is. Values close to 90 degrees show an ‘oblong’ flock shape.

#### Turning direction

Members of both real and simulated flocks were categorised based on their turning motion relative to the headings of the flock and the predator (Fig 2A). Following the definitions of Sankey *et al*. [24], when the direction away from the predator’s heading is also the direction away from the average heading of the flock, we will refer to the individual as being ‘in-conflict’ (between turning to either escape or align with the flock). In a non-conflict scenario, an individual needs to turn towards the flock’s heading in order to escape. According to these categories, an individual may actually turn as follows:

1. in a ‘conflict’ scenario:

a. towards the flock and the predator (potentially staying in the flock and risk getting caught),
b. away from the flock and the predator (potentially splitting-off and escaping);
2. in a non-conflict scenario:

a. towards the flock and away from the predator (potentially staying in the flock and escaping),
b. away from the flock and towards the predator (potentially splitting-off and risk getting caught).

**Fig 2.**
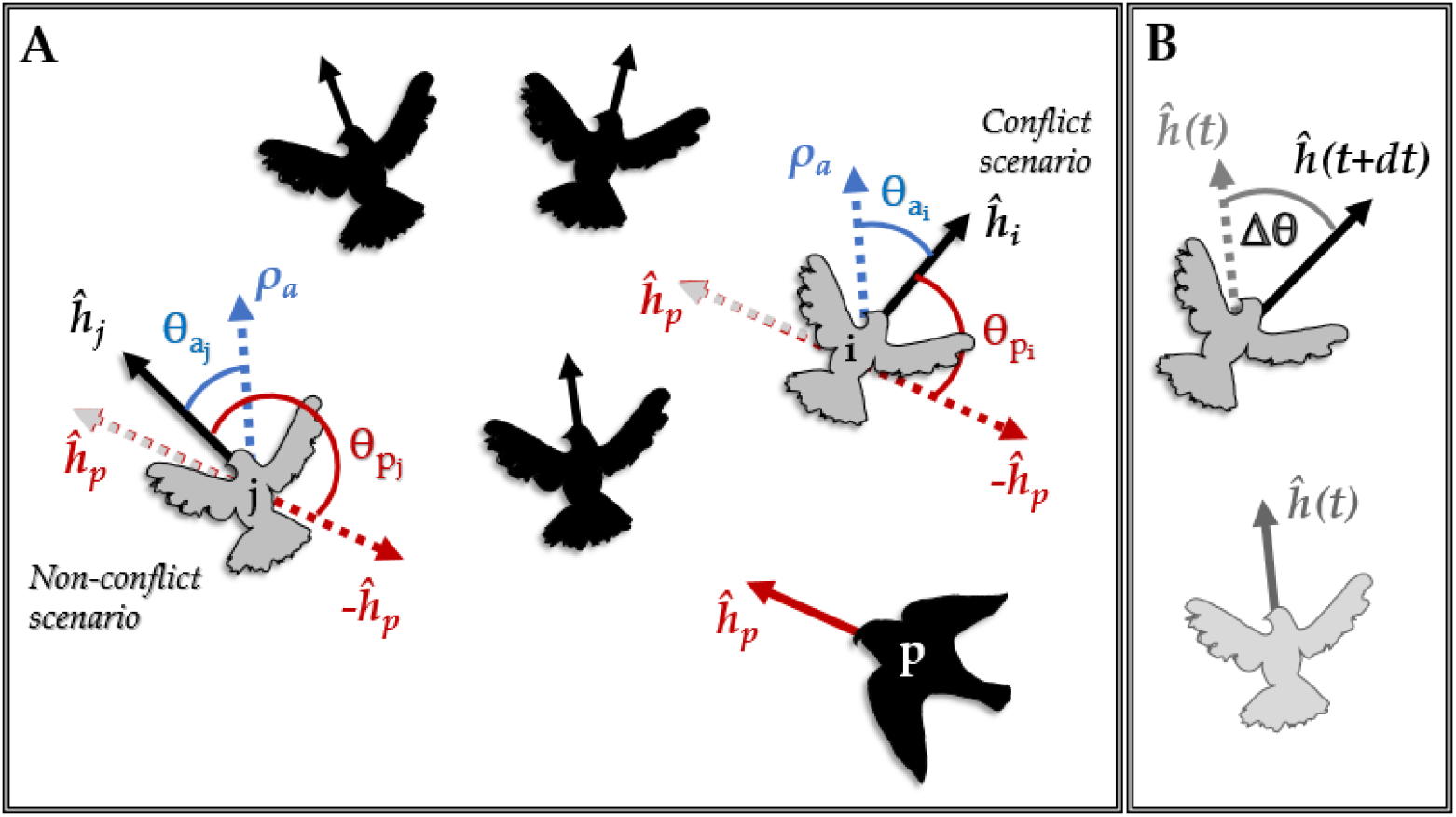
Definition of turning direction. (A) Prey individuals *i* and *j* are members of a flock with average heading *ρ_α_* and face different conditions to escape the predator *p*. The headings (unit vectors) of the individuals are represented by *ĥ*. The red angles *θ_p_* are the turns away from the heading of the predator and the blue angles *θ_c_*, the turns towards the heading of the flock. Based on the relative heading of individuals *i* and *j* to the headings of the flock and the predator, *j* needs to escape while aligning with its flock-mates, while *i* is in-conflict, since it needs to turn away from the flock in order to escape. (B) The change of heading of an individual between consecutive time steps (*dt*) is represented by *Δθ*. Its sign shows whether the individual turned towards the flock (same sign with *θ_c_*), away from the predator (same sign with *θ_p_*), towards neither or both (depending on its escape conditions).

We identified the turning direction of each flock member in respect to the predator by comparing the signs of a series of angles relative to the focal individuals heading (Fig 2A). Firstly, the direction ‘towards the flock’ is indicated by the sign of the angle *θ_a_*, the one between the focal individual’s heading and the average heading of the flock (*ρ_c_*). Secondly, the direction away from the predator (*θ_p_*) is shown by the sign of the angle between the prey’s heading and the vector (—*ĥ_p_*) opposite to the predator’s heading. Lastly, the direction of an individual’s motion is estimated as the sign of the angle between its heading at consecutive integration time-steps (Δ*θ*(*t* + *dt*), Fig 2B). After comparing these three directions, we categorised each individual’s motion in the above mentioned categories (1a, 1b, 2a, 2b).

As in the analysis of Sankey *et al*. [24], we split our data into 10-meter clusters of predator-prey distance (from 0 to 60 m). We calculated the frequency of each turning category at each cluster across all real and simulated trajectories. We refer to the frequency of turns away from the predator (1b, 2b) as ‘escape frequency’.

#### Distance dependency

We analyzed the simulated data focusing on how several variables scale with decreasing distance to the predator. Specifically, we calculated for each predator-prey distance cluster:

1. the angle (*α_pi_*) between the headings of the predator-agent (*ĥ_p_*) and each prey-agent (*ĥ_i_*) at each time step.
2. the frequency that pigeon-agents change their escape direction across all simulations:

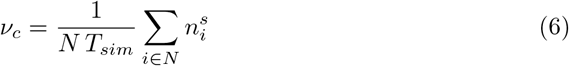

where *N* is the total number of pigeon-agents in all simulations, *T_sim_* the total simulation time, and 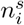 the number of occurrences that the escape direction of agent *i* changed between time steps (*θ_pi_* (*t*) = *θ_pi_* (*t* + 1)).
3. the consensus in escape direction in each flock at every time step:

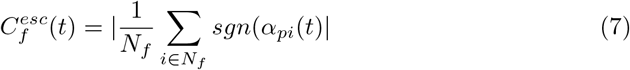

where *N_f_* is the number of individuals in the flock *f*, and *sgn*(*α_pi_*(*t*)) the sign of the angle between the headings of the predator and individual *i* at time *t*. Values close to 1 show that most flock-members have the same escape direction, whereas 0 indicates that half of the flock needs to turn to the right to escape and the other half to the left.

#### Self-organised dynamics

To inspect the interplay of coordination and predator-avoidance during collective escape in our model, we examined the effect of alignment and centroid-attraction on the motion of each individual. Specifically, we collected the weighted steering forces (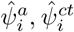, Eq 2) of each flock-member in its own coordinate system:

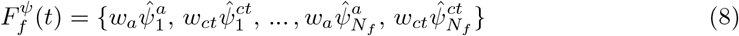

where 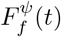 is the set of the coordination forces acting on all *N_f_* focal individuals of flock *f*. We used these sets to create density maps of coordination effect during a predator’s attack on a flock.

### Software

Our computational model was built in C++ 17. Graphics rendering was implemented in OpenGL on MS Windows^®^ OS [34]. The calculations of bearing angle, flock shape, predator-prey distance, and the determination of conflict scenarios for the simulated data were performed in C++. All other analyses of empirical and simulated data were performed in R (version 3.6). All result plots were made in ‘*ggplot2*’ [35].

## Results

### Characteristics of a flock of homing pigeons

We analyzed tracks of homing pigeons in flocks, initiating their homing flight or being attacked by a robotic falcon (*N* = 43). To characterize what comprise a flock of pigeons, we constructed for each flock the distribution of individual speed, nearest neighbor distance, and calculated for each focal individual the relative position of all its neighbors (distance and bearing angle) (Fig 3 A1-C1). We found large differences in these distributions among flocks (S1 Fig). The bearing angle to each nearest neighbor (S2 Fig) is uniform distributed (Kolmogorov-Smirnov test, comparison with a uniform distribution, D = 0.08, p-value = 0.58). The shape of each flock varies from oblong to elongated, according to our calculation of the angle between the flock’s heading and the shortest axis of the minimum-area bounding box surrounding all members (with an average angle of 41 ± 23.5 degrees). In total, in a flock of homing pigeons, an individual flies with an average speed of 18 m/s and have its nearest neighbor positioned anywhere around it (bearing angle from −180 to 180 degrees) at a distance of 1.3 ± 1.8 meters (mean and standard deviation).

**Fig 3.**
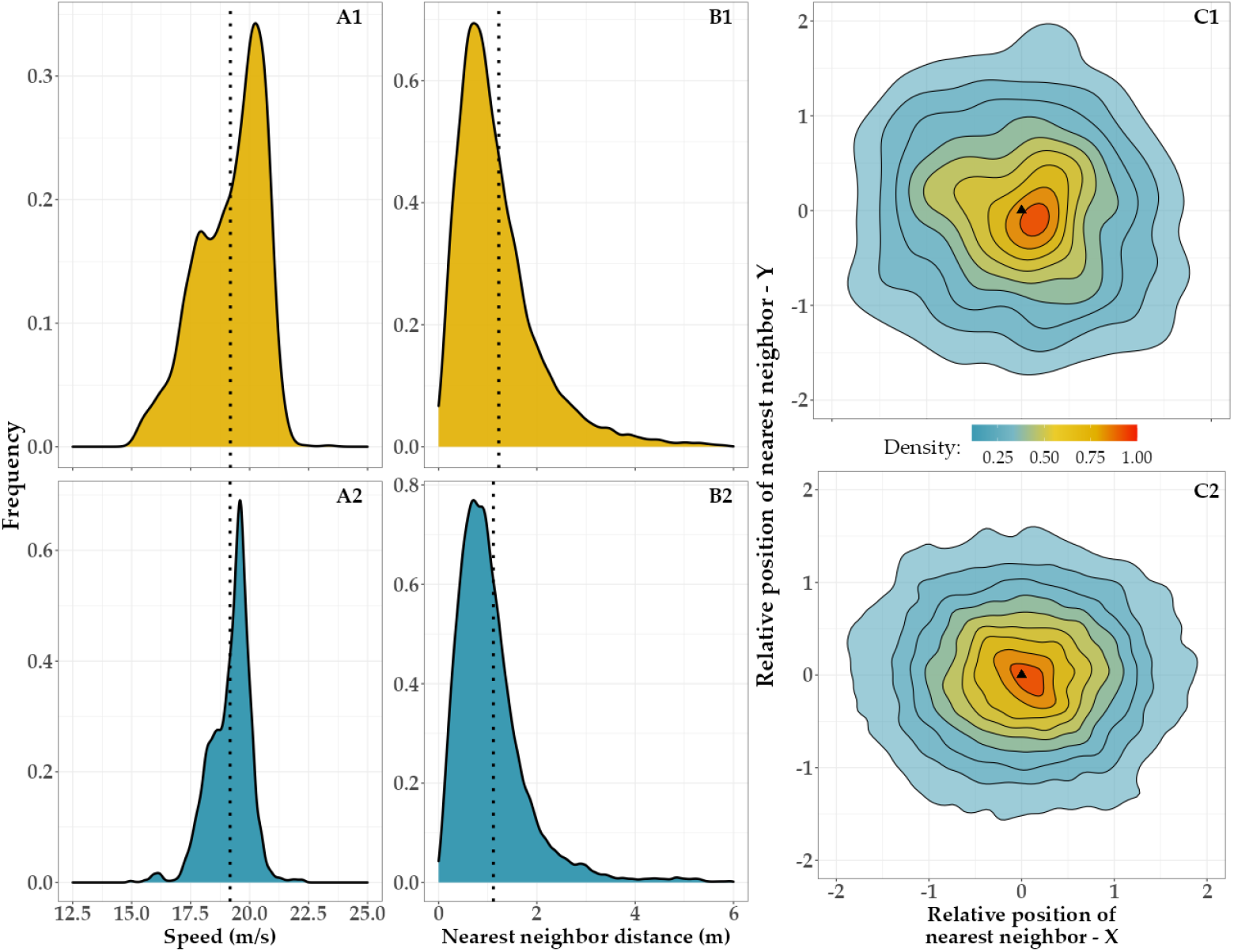
Comparison of real and simulated flocks of pigeons. Data from real (top row) and simulated (bottom row) flocks of 8 pigeons. (A-B) The vertical dotted lines show the mean of each distribution. (A) The distributions of individual speed throughout a flight of real pigeons (A1) and a simulation (A2). (B) The distributions of nearest-neighbor distance throughout a flight (B1) and a simulation (B2). (C) Density of the nearest-neighbor positions in the coordinate system of each focal individual, based on the bearing angle and distance to the nearest neighbor (*m*). Estimated for a real flight (C1) and a simulation (C2). The triangle represents the position of the focal individual heading to the north direction.

### Simulated flocks of pigeons

Before inspecting our hypothesis, we established whether the behavioral rules of our model result in pigeon-like flocks [17]. To develop our model, we adjusted the relative importance of the coordination rules (alignment, attraction, avoidance) among individuals according to empirical data of pigeons. For each pigeon-agent, we modeled strong alignment with the average heading of its 7 closest neighbors, and weak attraction to their center of mass. As to attraction, we also included an accelerating mechanism, as found by Pettit *et al*. 2013 [15]. While individuals accelerate to stay close to their flock mates, an additional force drags them back to their personalised cruise speed [21].

We measured the flock characteristics described above (distribution of speed, nearest neighbor distance and relative position of neighbors) in our simulated tracks. Due to the large variability in measurements among real flocks, the cumulative distribution across flights is not representative of individual flocks (S1 Fig). Hence, we compared the simulated data with tracks of a single flock and validated that our computational model closely resembles flocks of pigeons (Fig 3).

### Turning-direction frequency under attack

We model pigeon-agents to avoid the heading of the predator rather than its position, according to the findings of Sankey *et al*. 2021 [24]. Specifically, we added a force perpendicular (in ±90 deg angle) to the heading of each agent, pushing them to turn away from the predator (while also coordinating with their neighbors). The strength of this force is constant and independent of the predator-prey distance. Only its direction (the forces’ sign) changes during an attack, based on the relative direction of the prey’s and predator’s headings. Our simulations show a pattern very similar to the one of the empirical data: the closer the predator, the higher the frequency of escape is (Chi-Square for Trend test, X-squared = 10.3, p-value = 0.001, Fig 4B), independent of our predator-agent’s strategy (‘lock-on’ and ‘chase’, S3 Fig).

**Fig 4.**
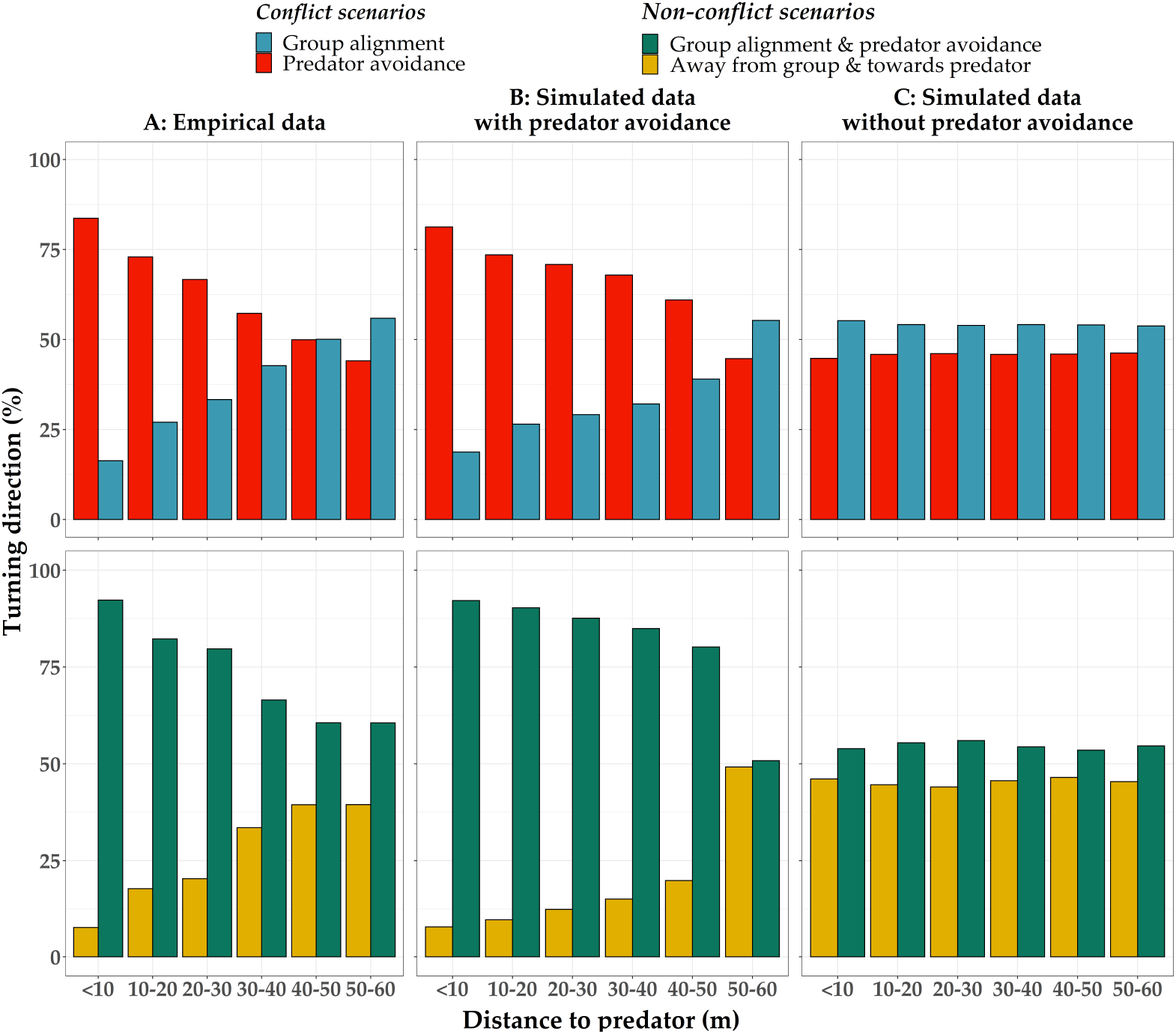
Turning direction frequencies of flock members. The percentage of turns towards the four turning directions at consecutive time-steps (for individuals under ‘conflict’ and ‘non-conflict scenarios) as a function of distance between them and the predator, across the empirical data and two simulation experiments. (A) Empirical data of Sankey *et al*. [24]. (B) Simulated data with predator avoidance that is independent of the distance between the predator and the prey individuals (modeled as in Fig1). (C) Data from control simulations where the prey does not react to the predator.

This shows that in our model, the pattern of collective escape that scales with predator-prey distance emerges from an individual behavior that does not take their distance to the predator into account (distance-independent avoidance). In simulations where pigeon-agents to not react to the predator (control), the frequency of escape remains constant (Chi-Square for Trend test, *x*^2^ = 0.03, p-value = .87, Fig 4C), as expected.

### Parameters scaling with predator-prey distance

We examined several aspects of our system to identify why pigeons turn more frequently away from the predator when they are closer to it (Fig 4). We focused on measurements that differ between simulations with and without predator-avoidance, to eliminate effects of mechanisms that are irrelevant to predation. When pigeon-agents do not react to the predator, the values of the chosen measurements show little or no variation with predator-prey distance (Fig 5. A1, B1, C1).

**Fig 5.**
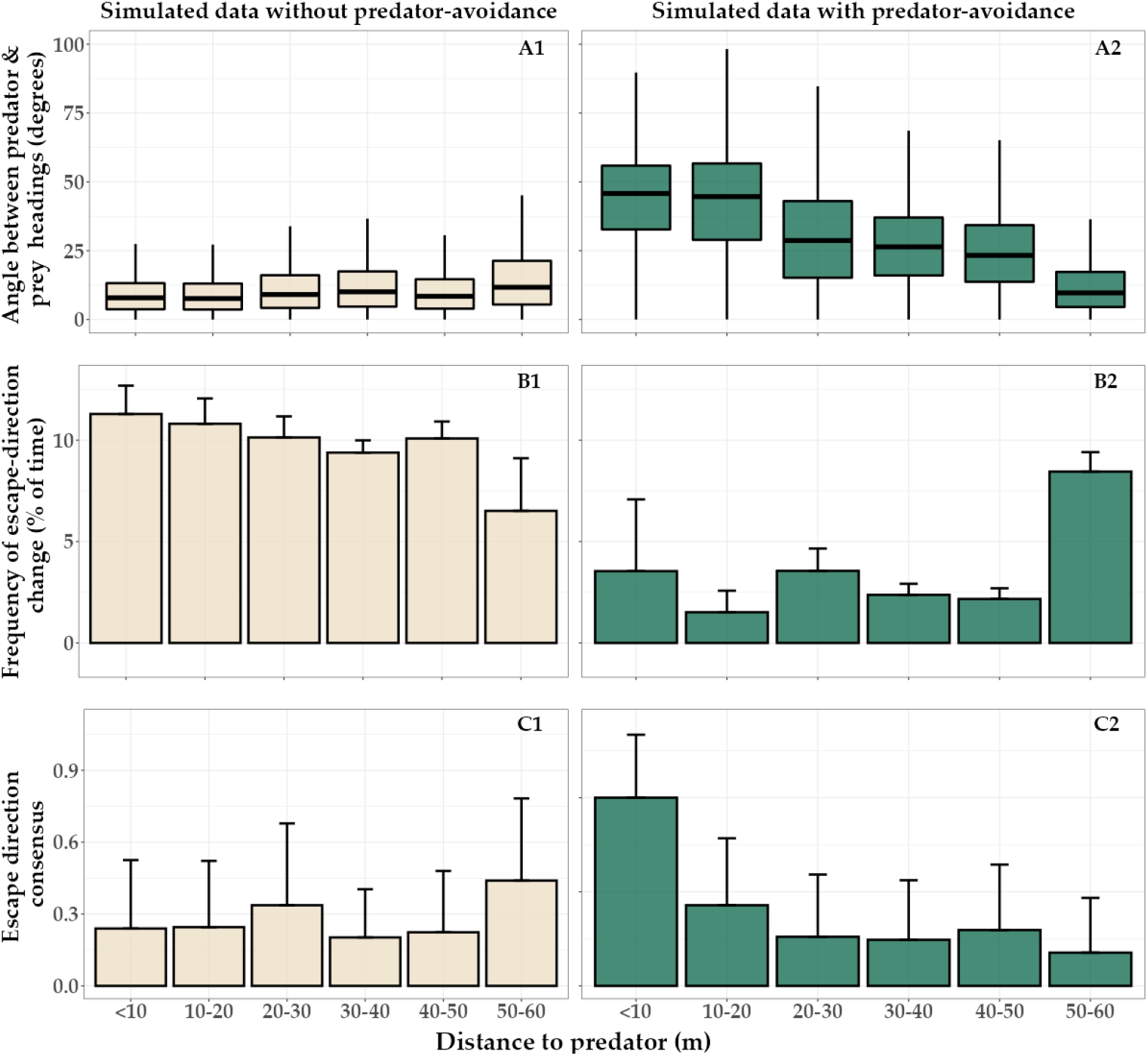
Distance-dependency in simulated flocks with and without predator avoidance. (A) The angle between the headings of the predator-agent and the pigeon-agents at each distance-cluster. The boxes include the 50% of the distribution and the horizontal line shows its median. When pigeon-agents react to the predator, the measurement increases showing that the flock is turning away from the predator. (B) The frequency that the escape-direction of each flock-member is changing direction (Eq 6). The height of each bar shows the mean value of all individuals per distance cluster and the error bar shows one standard deviation above this mean. The escape direction remains more stable when the flock is turning away from the predator. (C) Consensus in escape direction across a flock at each sampling point (Eq 7). More flock-members have the same escape direction closer to the predator in simulations with predator-avoidance.

Figure 5 A2 visualises the progression of a collective turn: the closer the predator, the larger the angle between its heading and the preys’ headings. By turning away from the heading of the predator, agents reinforce the predator-avoidance force to be towards the same direction (relative to their headings) across consecutive time steps. As a result, the frequency with which the predator-avoidance force changes direction decreases with decreasing distance to the predator (Fig 5. B2). When the predator is very close, the escape direction across the flock is almost constant (since the flock is already performing a collective turn) and the escape direction has the same sign for all individuals. From this mechanism, the consensus in escape direction across the group increases at closer distances to the predator (Fig 5. C2).

### The role of self-organization

By combining our results, we form a theory that this pattern emerges by self-organisation from the coordination among individuals during collective turning.

From the measurements that scale with predator-prey distance (Fig 5), we concluded that the progression of the collective turn is necessary for this pattern to emerge. Hysteresis [5] may play a role in the identified pattern on turning direction; the state of the flock at a specific distance to the predator has an effect on the pattern of the next state (at a closer distance). This was supported by our additional experiment where we positioned the predator close to the flock at initialisation with a speed lower than the flock’s. As the predator is drifting backwards (in an almost opposite motion to the one of its normal attack), the distance-dependent pattern does not arise (S3 Fig). Specifically, the turning-direction frequency does not differ among the clusters (< 10 to 50 meters) of predator-prey distance (*p* — *value* = 0.171, *X*^2^ = 1.875), in contrast to the effect we see in our main simulations (*p* — *value* < 0.005, *X*^2^ = 10.321).

To take a closer look at coordination among flock members during a predator attack, we extracted the effect of the coordination forces of centroid-attraction and alignment on the turning motion of our pigeon-agents throughout the simulations. In Figure 6, we see the details of coordination and the resulted collective motion during an attack. The simulated tracks show two consecutive collective turns of the flock as a reaction to the predator (Fig 6A). At intermediate distances (20-30 meters) to the predator, increased consensus in escape direction (Fig 5 C2) is causing the alignment and cohesion force to both pull individuals away from the predator into a turn. We describe the process below, referring to the simulated data of the single attack shown in Figure 6. We ‘ll classify the agents that are at the side edges of the flock, based on their relative position to the flock’s direction of turning, into ‘inner-edge’ (e.g. on the left side of the flock in a left escape-turn) and ‘outer-edge’ (on the opposite side).

**Fig 6.**
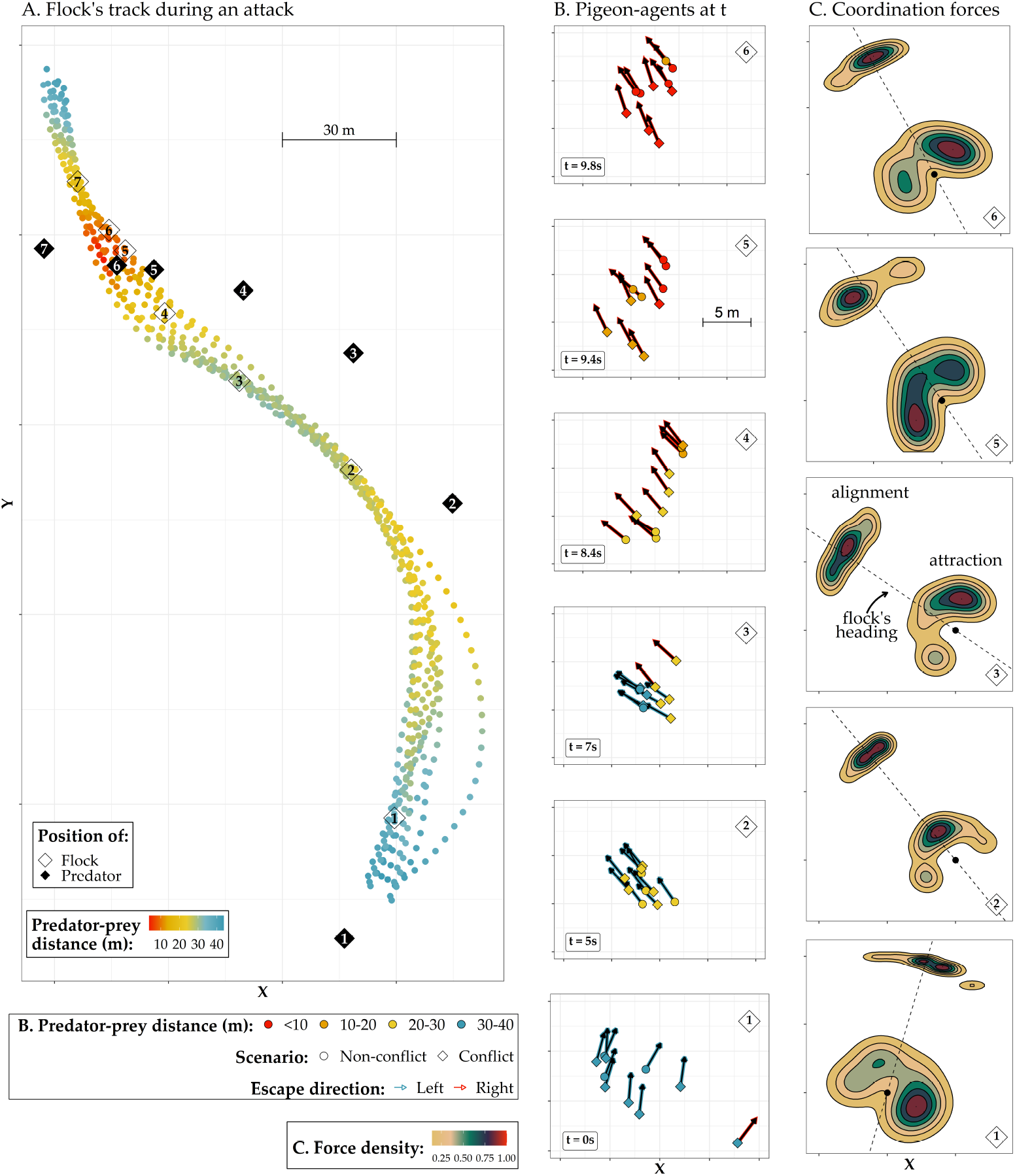
Progression of a collective escape. (A) The tracks of a simulated flock of 10 pigeons under attack. For simplicity, we present the part of the track when the predator is within 40 meters distance from the flock (excluding larger distances shown in previous plots). The points represent the position of single pigeons per time step of 0.2 seconds. The filled black rhombi show the position of the predator at 7 discrete time points. The numbers represent the link between the position of the predator and prey in time. (B) Positions of the pigeon-agents at time points 1 to 6 (0 to 9.8 seconds). Their color shows the distance to the predator of each individual (according to the clusters of Fig 4). Arrows represent the heading of each agent. The shadow of the arrows shows the escape direction of each agent at that time point. (C) The effect of centroid-attraction and alignment forces across the flock during the respective time points (density map of 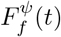, Eq 8). The point represents the position of all individuals in their local reference frame and the dotted line shows the average heading of the flock at that time point.

#### Predator’s behavior

As the predator is approaching, the flock is continuously turning away from its heading. We observe the predator getting closer to the flock after a change in the direction of its attack (as seen in Fig 6 A1 to A2), by crossing the flock’s path from the side (Fig 6 A5 to A6).

#### Flock shape

The shape of the flock has an important effect on the dynamics of a collective turn. In the beginning of a turn, the flock has a relatively wide shape (Fig 6 B1, B4). Exiting a turn, the flock becomes oblong (Fig 6 B2). In this shape, the centroid-attraction is in the direction of the flock’s heading (Fig 6 C2). Simultaneously, the alignment-force acts also closely around the flock’s heading (Fig 6 C2); the flock is very polarised. When the predator catches up and approaches from the opposite direction, the inner-edge individuals switch their escape direction first (Fig 6 B3). By starting to turn, they change the shape of the flock to wide (Fig 6 B3, B4) and move the effect of coordination forces (center of the flock and average heading) towards the escape direction (Fig 6 C3). With the coordination forces acting in the same direction as escape, the whole flock enters the turn (Fig 6 B4, B5). When this happens, the center of the flock re-positions to the opposite direction of escape (Fig 6 C5). The individuals that have started the turn (inner-edge) are now more attracted to the non-escape direction (Fig 6 C5). Outer-edge individuals (on the opposite side of the flock’s center) are still moving inwards (Fig 6 B5). The flock is becoming more oblong and approaches the exit of the turn (Fig 6 B6), where alignment and coherence will again have a subtle effect on the individuals’ turning (by acting around the flock’s heading, Fig 6 A7, as A-C2).

#### Progression of turn

At one time-point, flock members may be in different distance-clusters to the predator (Fig 6 B3-6). The individuals that are closer to it, at the inner-edge of the group, will establish the common escape direction first (Fig 6 B3, red shaded arrows). These individuals (‘inner edge’) make the flock shape wider and move the center of the flock and the average alignment towards the escape direction, initiating the turn (Fig 6 A3). Since the turn propagates from the inner-edge individuals, the outer-edge individuals (on the outside of the flock) have a delay in starting the escape turn (Fig 6 A4, B4). At this point, outer-edge individuals are not in conflict, since their escape direction matches the average alignment of the flock. Further in the progression of the turn, with more individuals turning to escape, the shape of the flock becomes more oblong. For the initiators of the turn (inner-edge), the center of the flock re-positions towards the non-escape direction (Fig 6 B5, C5). For the outer-edge individuals, the center of the flock is in the direction of escape (Fig 6 B5, C5). By sharply turning towards the escape direction to catch up with the flock that started turning earlier (Fig 6 A4-5), they are in-conflict, since for them alignment acts in the opposite direction (Fig 6 B5-6, C5-6). Reaching the exit of the turn, alignment acts close to individual’s headings and attraction switches back, in accordance with the escape direction for most flock members (Fig 6 C6). This makes the flock to continue the turn until the elongated shape where attraction and alignment are around the agent’s headings (Fig 6 A7, as in A-C2)

#### The role of ‘in-conflict’ individuals

During an escape turn of the flock, different individuals are ‘in-conflict’ between keeping up with the flock’s heading or avoiding the predator. At the beginning, inner-edge individuals are in-conflict, resulting in them initiating the turn (Fig 6 B3). When the predator gets closer, and further in the progression of the turn, the outer-edge individuals (the last to start turning) are in-conflict. This results in cohesion acting in accordance to escape. The observed increase in escape with predator-prey distance may be caused by alignment and attraction acting in opposite directions for individuals in-conflict close to the predator (Fig 7). By the direction change of the coordination forces and the change in the conditions that each individual is under (depending on their relative positions in the flock) during the turn, the flock manages to stay together, while different parts are ‘in-conflict’.

**Fig 7.**
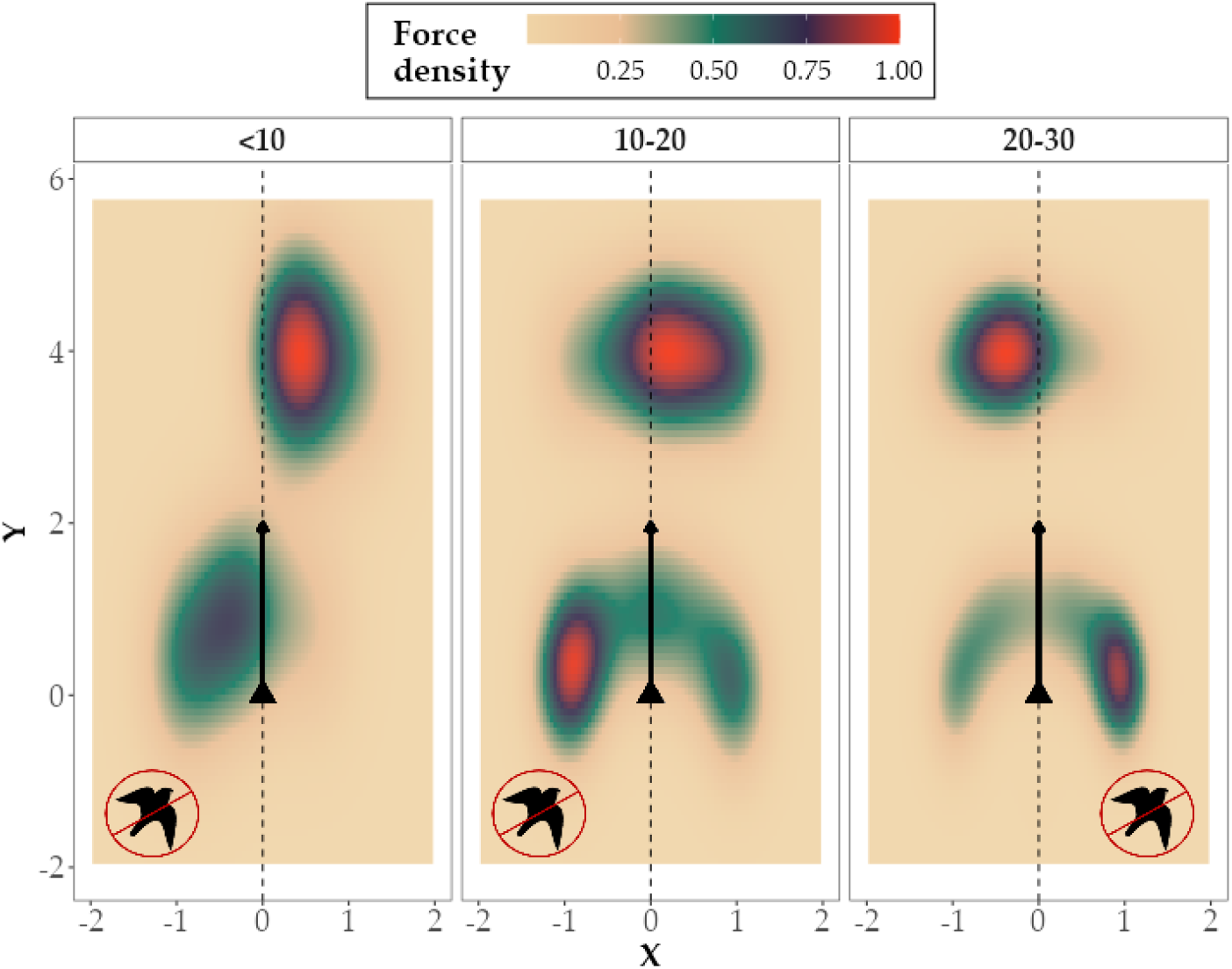
Effect of coordination forces on ‘in-conflict’ flock-members within 30 meters to the predator in *HoPE*. The density of turning-attraction (low center) and alignment (high center) forces (Eq 8) acting on the coordinate system of pigeon-agents that are in-conflict during the pursuit sequence shown in Fig 6. The triangle represents the position of each focal individual and the dotted line and arrow its heading. The predator sign represents the turning direction of predator avoidance. Alignment and centroid-attraction have opposite effects on turning direction relative to the agents’ headings and predator-avoidance is mostly in accordance with the centorid-attraction direction.

#### Mechanism’s summary

In total, at the beginning of the turn, in-conflict individuals are the ones initiating the turn due to the predator avoidance force. With the turn progressing, the flock shape becomes wide and by the in-conflict individuals having the center of the flock on their escape direction, cohesion acts in favour of escape. Towards the end of the turn, the initiators are moving more inwards, while the outer-edge individuals are the ones in-conflict, catching up with the escape turn.

In other words, when in-conflict at large distances (the beginning of the turn), only the predator-avoidance force acts in accordance with the escape direction, while the flock is oblong and polarised (thus coordination forces have small effect on turning). When the predator is closer (in the progression of the turn), attraction and alignment are in opposition. Attraction to the center of the flock is in accordance with the escape direction and the individuals turn more towards it.

## Discussion

Computational models based on self-organization have helped to unravel, in several group living organisms, what behavioral rules underlie complex collective phenomena [1, 5, 12, 19]. In the present study, we use such a computational model to explain how pigeons in flocks increase their frequency of turning to escape the predator, the closer they get to it. This distance-dependent pattern was initially hypothesized to reflect that prey rather turn away from the predator than align with their flock’s heading when threat increased [24]. We show in our model that this pattern emerges as a side-effect, without a distance-dependent rule for predator avoidance at the individual level. This implies that individuals do not need to prioritize escape over coordination to increase their escape frequency when the predator is close.

In reality, this mechanism may spare prey the cognitive effort [36, 37] of keeping track of the predator’s position during coordinated motion. We showed that consensus over an escape direction increases when the predator gets close, in a quorum-like response [38]. By individuals reacting only to the heading of the predator relative to theirs, the flock turns away from the predator’s direction of attack more as the predator gets closer.

The distance-dependent pattern emerged through self-organization from a combination of processes. First, a distance-independent tendency to turn away from the predator, leading the group into a collective turn. Secondly, the fact that centroid attraction acts in the direction of escape that is opposite to that of alignment when the flock is closer to the predator. Thirdly, through short-term hysteresis [5]. During the progression of a collective turn, as the predator approaches, the past state of the flock (shape and relative positions of flock members) affects its next state. Both, flock shape and internal structure, relate to what each individual experiences in terms of predator threat (angle of attack and escape direction) and social coordination (deviation from the flock’s heading and relative position to the centroid) and both affect the propagation of information concerning changes in heading through the group [39]. To our knowledge, the effect of hysteresis is new in the context of collective escape.

Predation is known to affect the coordination within groups of prey [39–41]. An example is a decrease in the minimum separation distance [40] and a potential increase in the number of interacting neighbors [41]. In fish, such changes often lead to increased group density [40], a pattern not seen in pigeons [24]. A decrease in minimum separation distance or increase in centroid-attraction in bird flocks may enhance the danger of collision while individuals turn away from the predator during flight. According to our results, a stronger tendency to align with flock members than to turn towards the flock’s center increases the prey’s escape frequency at shorter distance to the predator while retaining flock cohesion during collective turns. We thus hypothesize that for small flocks that turn away from their predator, increased alignment rather than decreased group density enhance chances of survival.

Whether prey escape by minding the predator’s position or its heading is unclear. In fish schools and insect swarms, individuals are supposed to avoid the position of the predator [42, 43]. Homing pigeons instead were observed to turn away from its heading [24]. In our simulations, we observed that with heading-avoidance a common escape direction is enforced among group-members supporting group cohesion during collective escape. In a previous model of fish schools, we see that when individuals turn away from the position of the predator, the group splits more frequently when the predator gets closer [7]. These avoidance strategies may be adaptive, depending on their effectiveness across species and ecological contexts. For instance, if prey is very maneuverable (i.e. fish and insects rather than birds [40, 44–47]) or subject to surprise attacks by their predator [46, 48], position avoidance may be more favorable.

The use of remotely-controlled predators [24, 25] helps to understand the relation between, on one hand, the coordination among group members and their rules of escape and, on the other hand, their collective escape behavior and locomotion specifics. By controlling the predator’s motion, the effect of different hunting strategies on the escape of the prey, both individually and collectively, can be tested. Additionally, collecting data on the order of different patterns of collective escape during a predator’s pursuit [22] can help reveal more effects of hysteresis on collective escape.

Our findings are relevant for collective escape by homing pigeons given that flocks in our model resemble those of real pigeons not only in their behavioral rules [15, 21, 24], but also in their emergent properties (e.g. distributions of speed and nearest neighbor distance) [17]. Similarly to previous species-specific models of collective behavior of different species [3, 12, 20], our model can be further used to study other aspects of flocking in pigeons, be extended to investigate evolutionary dynamics of collective escape [49], or be adjusted to other bird species. The increasing availability of quantitative data of collective behavior can, and should, further support the development of models around specific species to help interpret empirical findings.

## Supporting information

**S1 Fig. Distributions of speed, nearest neighbor distance, and shape of flocks of homing pigeons.** Each histogram shows the distribution of one flock during a control flight (based on the data of [24]). The bottom row shows the overall distribution across flights.

**S2 Fig. Distributions of bearing angle to nearest neighbor in flocks of homing pigeons.** The overlapping histograms show the distribution of one flock during a control flight (based on the data of [24]). The bottom row shows the overall distribution across flights.

**S3 Fig. Turning direction from all alternative simulation experiments with different predator strategy**. The default strategy is the ‘chase closest prey’.

**S4 Fig. Testing the accuracy of the topological-range estimation method of Sankey *et al*. 2021 [24]**. Their method is based on simple linear models between, on one hand, the turn that each individual performs during consecutive sampling points and, on the other hand, the turning angles for centroid-attraction and alignment. Angles based on all possible topological ranges for each flock size were tested. The linear model with the most explanatory power was througt to include the ‘real’ topological range. For the exact method description see [24]. Given that this method is not well established, we tested its performance on our simulated datasets. Specifically, we applied this method on data from simulations in which we vary separately the topological range for alignment and centroid-attraction (from 1 to all neighbors) for three flock sizes. We run 5 repetitions of each simulation with all unique combinations of topological range for alignment and centroid-attraction per flock size. The deviation index shows the deviation of the topological estimate of the linear-models method from the real value of topological range as parameterised in the model (shown on the x-axis), divided by the maximum possible deviation for each topological range and flock size (n-2, e.g. the maximum deviation for a flock of 8 individuals is 6 neighbors, when the true value is 7 and the estimate is 1 or vice-versa, giving a deviation index of 1). Values close to 0 show a good performance of the linear-model method. The method seems to lose accuracy when agents align with many topological neighbors and when they are attracted to the centroid of a few. Each point shows the mean deviation index of all simulations with the respective topological range and the error bars the standard error.

