## Supplementary figures and images for "Self-organization of collective escape in pigeon flocks"

### S1_Fig.tif

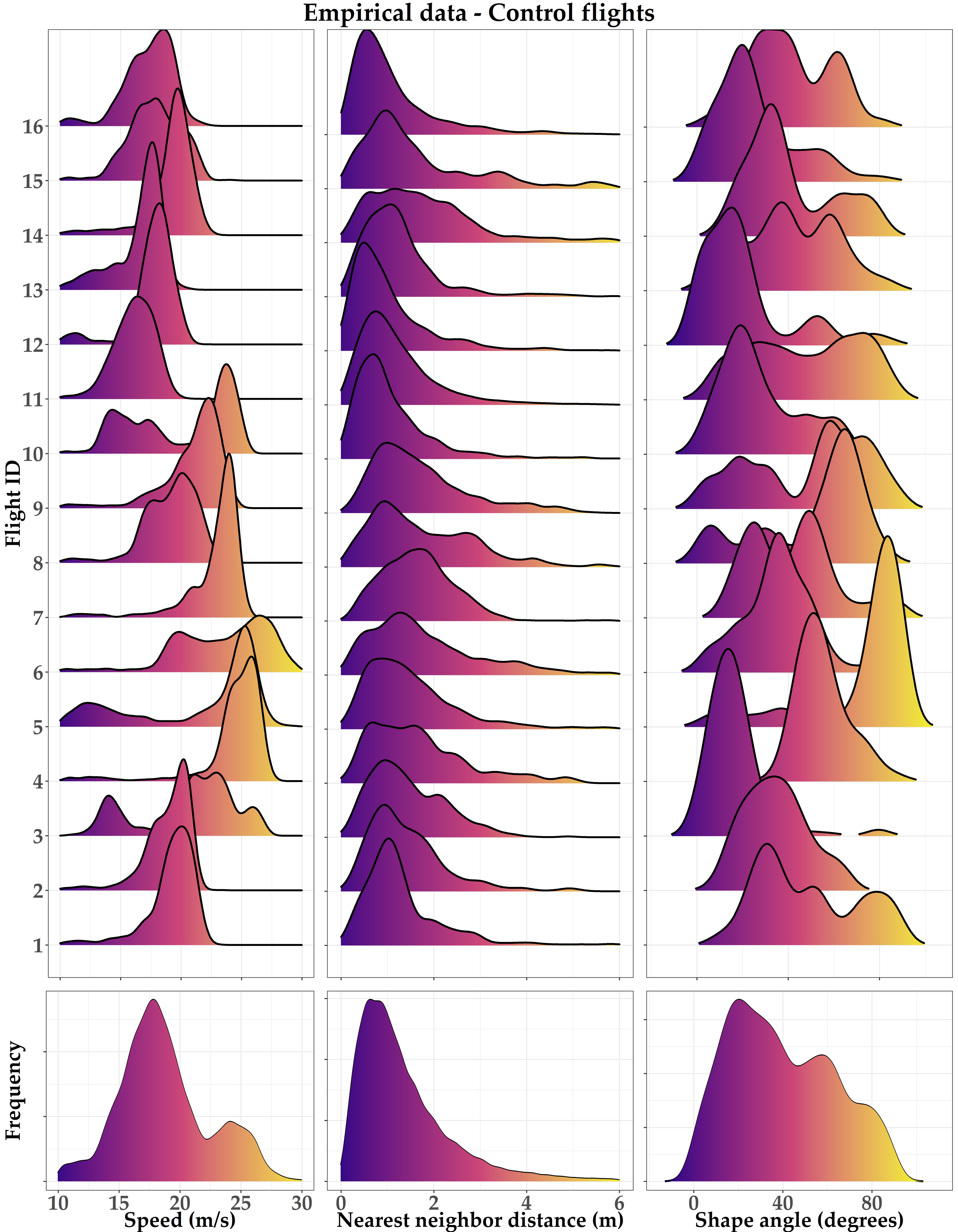

### S2_Fig.tif

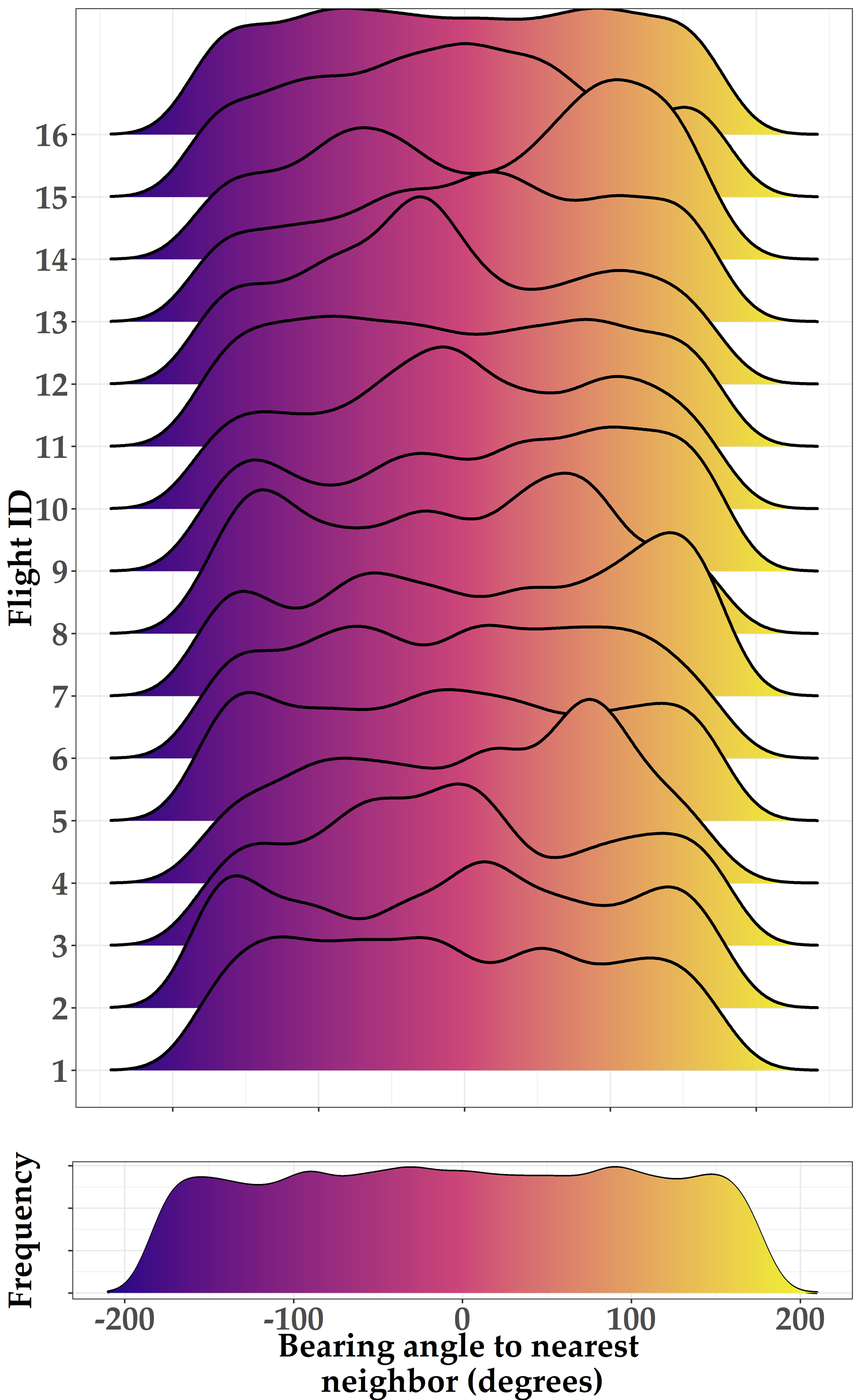

### S3_Fig.tif

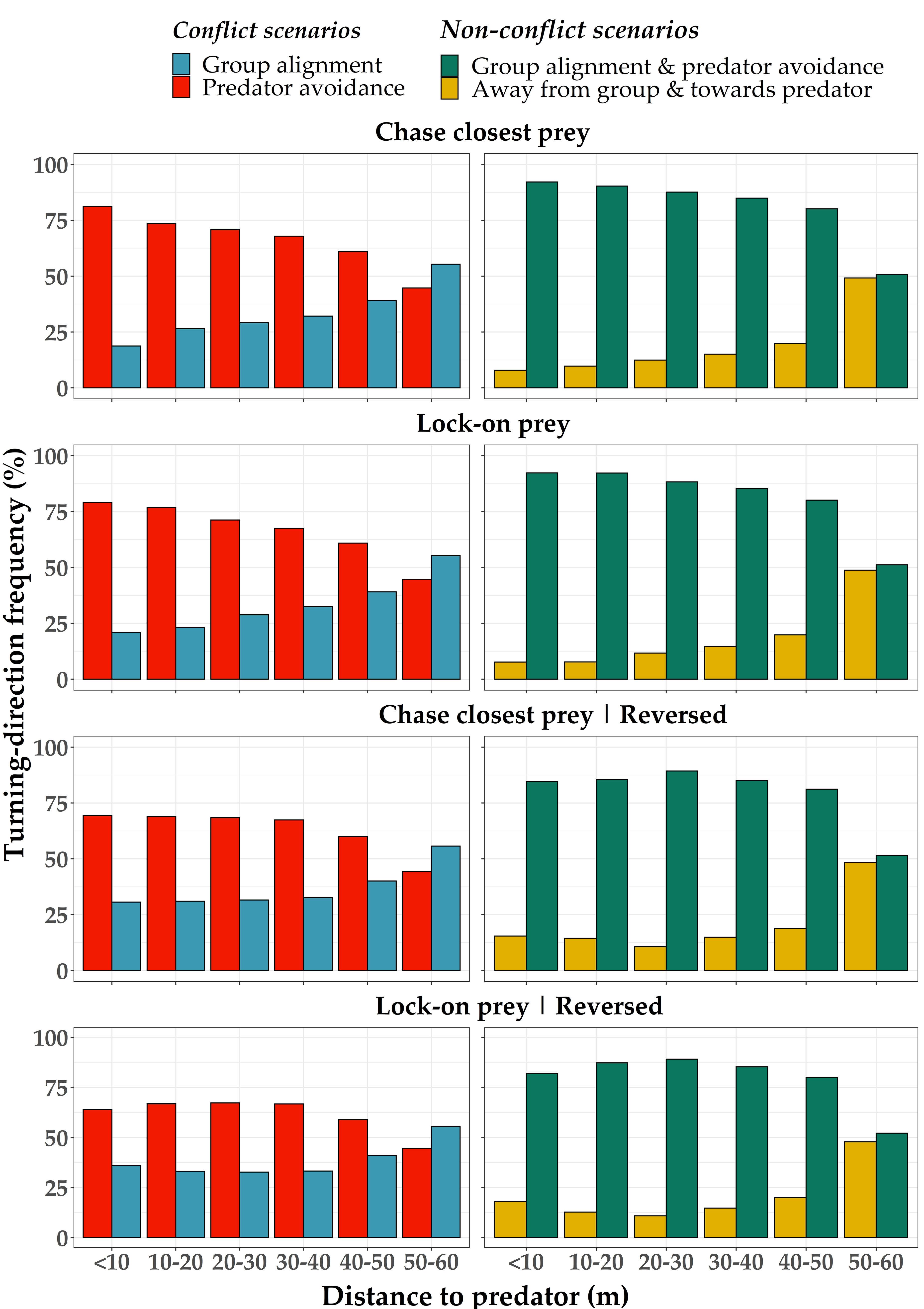

### S4_Fig.tif

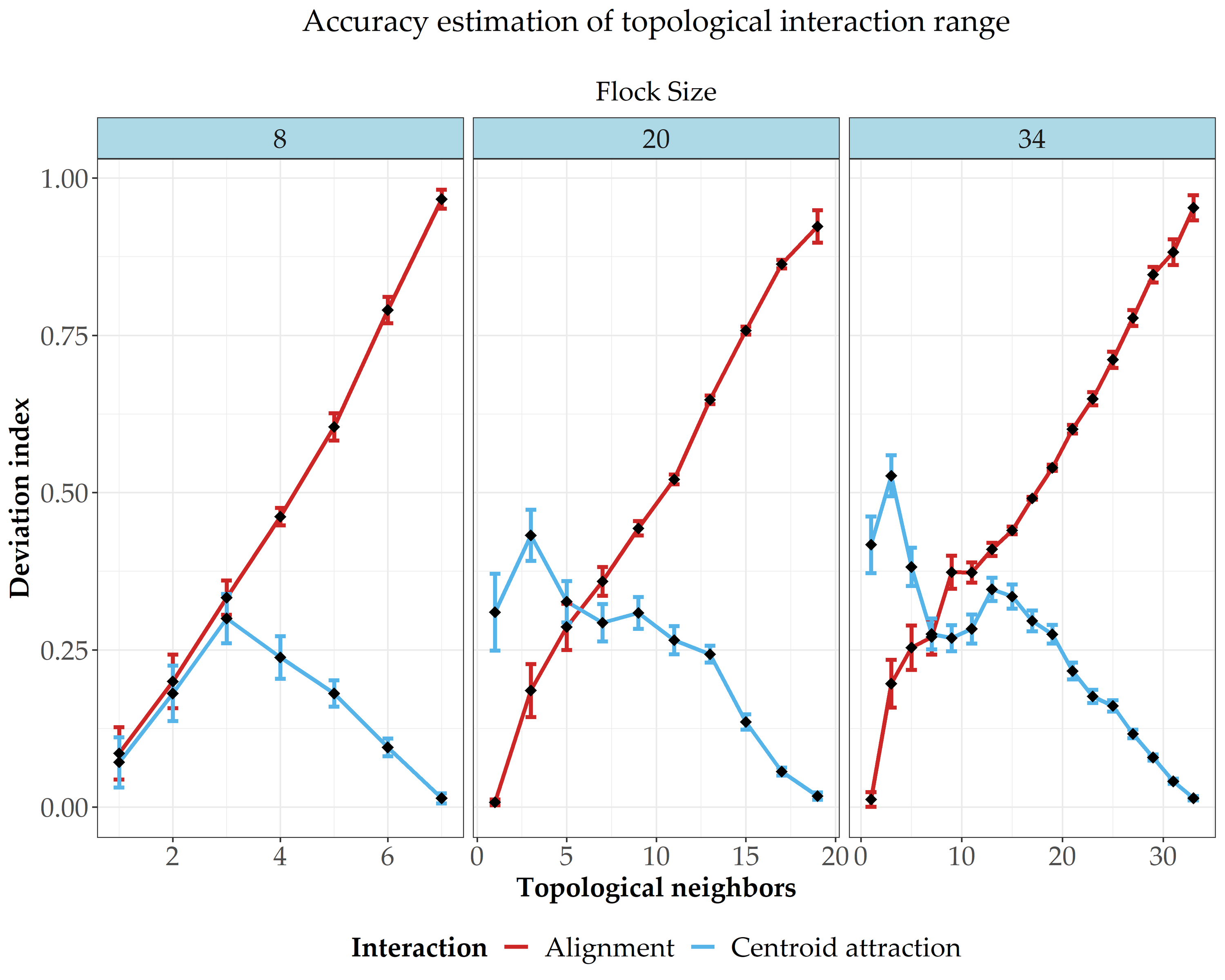
